# Spatial resource heterogeneity creates local hotspots of evolutionary potential

**DOI:** 10.1101/149302

**Authors:** Emily Dolson, Charles Ofria

## Abstract

Do local conditions influence evolution’s ability to produce new traits? Biological data demonstrate that evolutionary processes can be profoundly influenced by local conditions. However, the evolution of novel traits has not been addressed in this context, owing in part to the challenges of performing the necessary experiments with natural organisms. We conduct *in silico* experiments with the Avida Digital Evolution Platform to address this question. We created eight different spatially heterogeneous environments and ran 100 replicates in each. Within each environment, we examined the distribution of locations where nine different focal traits first evolved. Using spatial statistics methods, we identified regions within each environment that had significantly elevated probabilities of containing the first organism with a given trait (i.e. hotspots of evolutionary potential). Having demonstrated the presence of many such hotspots, we explored three potential mechanisms that could drive the formation of these patterns: proximity of specific resources, variation in local diversity, and variation in the sequence of locations the members of an evolutionary lineage occupy. Resource proximity and local diversity appear to have minimal explanatory power. Lineage paths through space, however, show some promising preliminary trends. If we can understand the processes that create evolutionary hotspots, we will be able to craft environments that are more effective at evolving targeted traits. This capability would be useful both to evolutionary computation, and to efforts to guide biological evolution.

## Introduction

Understanding the evolutionary processes that facilitate the evolution of complex traits is at the heart of many open questions in artificial life, evolutionary computation, and evolutionary biology. Numerous studies have suggested that spatially complex environments can drive the evolution of complex traits (Doebeli and Dieckmann, 2003; Wagner et al., 2013; Dolson et al., 2017), but it is unknown whether this result is a global pattern that is only visible at the scale of the entire environment, or whether it is driven by local spatial dynamics. If sub-regions exist that disproportionately promote the evolution of specific adaptive traits (*evolutionary hotspots*), they would be a valuable phenomenon to understand. Understanding what factors produce evolutionary hotspots should help us design environments that are more likely to evolve complex traits. This ability would also enable us to build more interesting artificial life systems, more effective evolutionary algorithms, and biological environments that promote desirable evolutionary outcomes.

Where should we expect to find evolutionary hotspots? Instinctively, we might expect a trait to arise only in regions where it is beneficial. However, selection can only favor traits once they appear, not promote that initial appearance, so other aspects of the environment must play a critical role. At the other extreme, hotspots seem unlikely in a homogeneous environment with no spatial structure. However, few environments in nature are truly homogeneous, and almost all have some form of spatial structure. Even among digital systems, once we consider the fact that any kind of interaction between neighboring individuals introduces spatial heterogeneity, few are truly spatially homogeneous. Thus, understanding if and how evolutionary potential varies across space could be important for understanding evolutionary dynamics in a variety of systems.

Why should we expect evolutionary hotspots to exist? To better understand this idea, it is useful to consider Wright‘s fitness landscape metaphor (Wright, 1932). At its simplest, a fitness landscape is a three-dimensional surface representing two traits (x and y) with varying values, and the fitness of an organism having a given combination of those trait values (z). When there are gradients in this space, we expect populations to climb to a fitness peak over time. Typically, populations will then get stuck at the peak of a local gradient, regardless of whether it is the highest location in the landscape. As such, the topography of the local fitness landscape plays a pivotal role in determining what traits will evolve in a population. A wide variety of factors can influence what part of a fitness landscape a population ends up in at a given point in time (Wilke et al., 2001). We predict that spatial variation across the environment is one such factor. A spatially heterogeneous environment can be conceptualized as a network of different fitness landscapes, one for each location in space. However, in a dynamic, interdependent ecological community, these fitness landscapes shift over time. Some local landscapes will tend to guide portions of the population toward regions of mutational space from which new traits are more easily accessible.

Of course, this network of fitness landscapes may not guide evolving populations in a consistent direction. While it is possible that landscapes could align to perfectly guide a lineage through the most challenging steps in the evolution of a complex trait, this condition seems unlikely. We might just as easily expect that the lack of a constant landscape will only serve to increase stochasticity, rather than having a predictable effect. Perhaps the fitness landscapes in adjacent locations even work at cross-purposes, with each guiding the population in opposite directions. Given this uncertainty, the primary goal of this paper will be to determine whether hotspots actually occur with sufficient frequency and strength to be worthy of further consideration. Addressing this question requires vast numbers of repetitions of experimental evolution in complex spatial environments. Conducting such an experiment in a wet lab system would not be worth this immense effort until additional theory has been developed. Once we have generated concrete hypotheses about what we would expect to see in biology, more targeted follow-up experiments in a wet lab system may be worthwhile. Here, we lay the groundwork for such theory by performing experiments in a digital system, where we will be able to precisely control the spatial distribution of environmental conditions and measure the drivers behind any spatial variation in evolutionary potential that we detect.

Here, we focus on a simple environment to avoid adding unnecessary complications to an already complex question. To this end, we use sessile, asexual organisms in an unchanging abiotic environment where different sets of traits are advantageous in different regions. Importantly, these traits are related to each other. This set-up is roughly analogous to a wide variety of scenarios within biology, ranging from bacteria surviving on a surface treated with a range of antibiotics (similar to (Baym et al., 2016)) to plants adapting to survive on different soil types within a region.

## Biological Background

Biologists have traditionally studied patterns of diversification rather than patterns of novel trait evolution (but see (Baym et al., 2016)). These processes have important distinctions from each other; diversification usually requires either physical separation of sub-populations or conflicting selection pressures, whereas novel trait evolution does not. Moreover, they are maximized by different conditions (Walker and Ofria, 2012). Nevertheless, diversification and novel trait evolution are closely related, as diversity is thought to increase evolutionary potential (Rouzic and Carlborg, 2008). The most well-studied biological pattern pointing to spatial variation in evolutionary processes is the latitudinal diversity gradient: a consistent pattern of high biodiversity near the tropics, and progressively less toward the poles (Hillebrand, 2004). While it remains impossible to perform controlled evolution experiments on a planetary-scale, progress has been made on untangling the drivers of this pattern. Most importantly for the purposes of this paper, evidence is mounting that diversification rates are higher in the tropics (Mittelbach and others, 2007). The drivers of this variation are still debated, but many hypothesized drivers are related to spatial dynamics within the environment, most notably the idea that the tropics may contain more environmental gradients, barriers to gene flow, and other spatial properties that promote speciation (Moritz et al., 2000; Doebeli and Dieckmann, 2003). Although these dynamics are more complex than those addressed here, particularly given that many rely on sexual reproduction, they suggest that spatial variation in evolutionary processes has broad relevance.

In related work, conservation biologists have sought to locate regions with an elevated intensity of evolutionary processes. This research stemmed from an increasing recognition that preserving the processes that generate diversity is at least as important as preserving existing diversity (Moritz, 2002; Ferrire et al., 2004). In accordance with this motivation, these studies tend to focus on small regions of land, making them more directly comparable to the research presented here. At a global scale, certain regions simply lack the genetic background for a given trait to have a chance of evolving (e.g. a novel heat tolerance mechanism seems unlikely to evolve in the arctic). Our question centers on more localized regions that are spatially heterogeneous with regard to the extent to which various related traits are useful. The results from conservation biology suggest that variation in evolutionary potential on such a local scale is plausible; even within relatively small regions, these studies have demonstrated substantial spatial variation in phylogenetic diversity (Forest et al., 2007; Vandergast et al., 2008).

Most of the processes that are hypothesized to be involved in the formation of hotspots of phylogenetic diversity are based on insights from simpler systems, most notably island groups. Islands represent clearly delineated spatial patterns and processes, making them an excellent context in which to build up insights about how eco-evolutionary dynamics will play out across space. For example, islands are known to promote diversification via adaptive radiation (Losos, 2010), and the formation of new islands within a system has substantial impacts on the assembly of ecological communities over evolutionary time (Gillespie, 2004). Island systems are also a particularly pronounced example of variation in spatial structure. Spatial population structure has a profound effect on how evolution will proceed; more structured populations (i.e. those with more constraints on where offspring end up relative to their parents) are generally more diverse and more likely to reach the highest peak in a fitness landscape (Tomassini, 2005; Nahum et al., 2015). The theory of island biogeography makes various shorter timescale predictions (MacArthur and Wilson, 1967). More recently, biologists have begun to translate theory developed for island systems to more continuous land masses, often by treating regions of a species’ preferred habitat as”islands” (also called patches) within a matrix of less suitable habitat (Forman, 1995). Many have argued this approach is likely an oversimplification (McGarigal et al., 2009; Franklin and Lindenmayer, 2009). A better understanding of the relationship between the dynamics observed in island systems and the dynamics that occur in more continuously varying environments will likely be important to understanding the spatial evolutionary dynamics that create evolutionary hotspots.

## Methods

### Study System

As previously discussed, addressing this question requires a system with a relatively complex abiotic fitness landscape. The Avida Digital Evolution Platform is an obvious choice for such a system (Ofria and Wilke, 2004). Not only does it easily allow for the creation of such fitness landscapes, it also includes a variety of tools for analyzing them. Avida is a world of self-replicating linear computer programs competing for space; when a program copies itself, it overwrites one of its 8 Moore neighbors. This is the only form of interaction between organisms. Mutations are introduced during the self-replication process, resulting in evolution via natural selection, with programs that can copy themselves faster having higher fitness. Improvements in fitness can either come from optimizations to the code of the genome (increasing its efficiency) or from gaining the ability to take in resources by performing boolean logic operations. Resources give programs extra CPU cycles, allowing them to copy themselves faster than competitors. Each of the nine two-input Boolean logic functions allows organisms to use a different resource, with more complex functions corresponding to more valuable resources. Here, we will use the ability to perform these functions as the traits of interest, and define organism phenotypes as the set of traits they possess.

Programs in Avida inhabit a 60 x 60 grid and offspring are placed in cells adjacent to their parents. Results presented here use non-toroidal (bounded) grids for ease of visualizing lineage trajectories (see 5), but otherwise identical experiments on a toroidal grids produced similar results. We create heterogeneous environments by placing different resources in each cell. For a previous experiment (Dolson et al., 2017), we created 500 environments by placing a random number of randomly-sized circular patches of resources in random locations across the environment. This technique produced a variety of environments while maintaining a correlation between the resources present in one cell and the resources present in its neighbors. For this study, we have arbitrarily selected eight environments for further study without examining them beforehand. In each environment, we performed 100 replicate runs of Avida. Each replicate was run for 100,000 updates (approximately 2,000 generations). We also performed 100 replicate runs of a homogeneous control condition where all resources were globally available. From these results, we extracted the coordinates of the grid cell in which each trait first appeared within each run. While a trait may evolve independently multiple times within a run of Avida, including these subsequent evolutions would introduce various biases into the dataset (especially when traits were lost and then re-evolved). Thus, we consider only the first organism ever to display a given trait within a replicate, before selection has an opportunity to play a role.

Our extracted (x,y) coordinates translate into 81 separate point patterns (one for each of the 9 traits across each of the 8 environments plus the control) (see Figure 1A): one for each of the nine different traits in each of the eight environments. Each pattern contains a maximum of 100 points. Many patterns contain fewer points, as it is not guaranteed that every trait will evolve in every replicate. Because each replicate is completely independent of each other replicate, the only relationship between points in the same pattern is that they evolved in the same environment.

**Figure 1:**
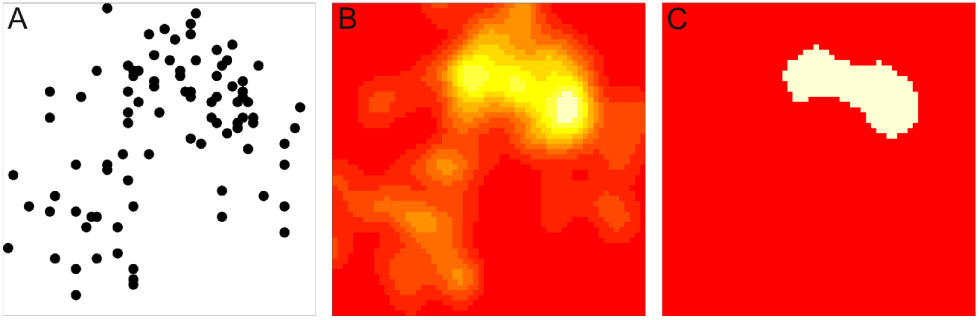
Hotspot location technique. A) An example point pattern, generated from 100 runs of Avida within a single environment. This pattern corresponds to the most complex trait, EQU. Each point represents the location in space of the first organism within one of the 100 runs that was able to perform EQU. B) Kernel intensity map generated from the point pattern in part A. C) Regions within that kernel intensity map that are substantially more intense than observed in the Monte Carlo simulations (i.e. the hotspot).

### Statistical Approach

To determine whether or not different regions have different evolutionary potential, we need to determine whether the point pattern is significantly different from complete spatial randomness. To make it possible to appropriately correct our statistics for multiple comparisons, we performed the test for spatial randomness in two steps: 1) a test to see whether each pattern, as a whole, was random, and 2) a follow-up test to determine which regions of non-random patterns had significantly more points than expected by chance.

For the overall randomness test, we calculated a test statistic for various distances, *r*. For each point in the pattern, we counted the number of other points falling within *r* distance units of it. We then averaged this number across all points to get our test statistic for that value of *r*. For each point pattern, we generated a range of expected results under the null hypothesis of complete spatial randomness by running 100 Monte Carlo simulations where we randomly placed the number of points in our actual data across an area of the same size. If the value of the test statistic calculated from our observed data was greater than the highest value from the Monte Carlo simulations for more than 5% of the r values, we concluded that the observed pattern was substantially more clumped than we would expect to see by chance (see Figure 2). A test statistic below the range of those observed for the Monte Carlo simulations would suggest that the observed data was more uniform than we would expect by chance; such a result would be unexpected for our experiments. Note for those familiar with spatial statistics: this statistical test is functionally equivalent to Ripley’s K function (Ripley, 1981). However, since Ripley’s K function assumes first-order spatial stationarity and our data is exclusively the result of first-order trends, we do not expect to see the same patterns traditionally observed in Ripley’s K function.

**Figure 2:**
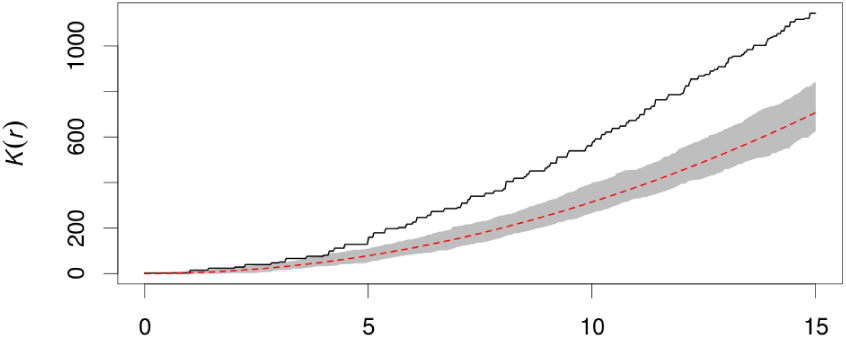
Results of performing the test for spatial clumping on the point pattern from Figure 1A. Observed values of the test statistic (black line) are well above the values observed in the Monte Carlo simulations (shaded area), indicating a clumping in this pattern. The red line indicates the theoretically derived expected value for a random pattern.

For those point patterns that were significantly more clumped than we would expect by chance, we performed a follow-up test to determine the location of the hot spot. We calculated a kernel intensity surface (essentially a heat map of point density across the environment) for each of these point patterns using the spatstat library (Baddeley et al., 2015) (see Figure 1B). Then, to determine whether the patterns within the data were significantly more intense than we expect from a random process, we performed a Monte Carlo hypothesis test with 100,000 simulations. We simulated patterns generated under the assumption of complete spatial randomness (our null hypothesis) by randomly selecting points from a uniform 60x60 lattice grid approximating the lattice of cells in Avida. For each point pattern, we selected the number of points observed in the original pattern. Kernel intensity values were created for each of these simulations. For each cell in the grid, we compared the experimentally-derived intensity to the distribution of simulated intensity values. Cells in which our observed value was higher than 99.999% of the simulated values were labeled as hotspots of evolutionary potential for a given trait. This significance threshold (*p* < .00001) was selected to ensure all results are significant after a sequential Bonferroni correction for multiple comparisons across all 3600 grid cells.

### Code and data availability

Avida is open source and freely available at https://github.com/devosoft/avida. Scripts used to generate environment files are available via Zenodo with DOI 10.5281/zenodo.162981. All statistical analyses were performed using the spatstat library, the qqtest library, and the R Statistical Computing Language (Baddeley et al., 2015; Oldford, 2016; R Core Development Team, 2013). Data and scripts used in this project are available at https://github.com/emilydolson/evo_in_space.

## Results and Discussion

### Existence of evolutionary hotspots

Out of the 81 point patterns, 43 differed significantly from random. All 43 of these non-random patterns showed significant clumping of points. For 40 of these 43 non-random patterns, the kernel density map had at least one region of higher intensity than we would expect to see by chance (hotspot) (see Figure 3). It is possible that the three patterns without hotspots instead had areas of significantly decreased evolutionary potential. Alternatively, these patterns may contain hotspots that were too weak to be detected by our highly conservative threshold for significance. Every test environment had hotspots for at least two traits, while none of the point patterns from the spatially homogeneous control displayed significant clumping, suggesting that these hotspots are a product of the spatially heterogeneous environment. These results provide clear evidence that hotspots of evolutionary potential not only exist but are relatively common; some regions of the environment promote the evolution of new traits to a greater extent than others.

**Figure 3:**
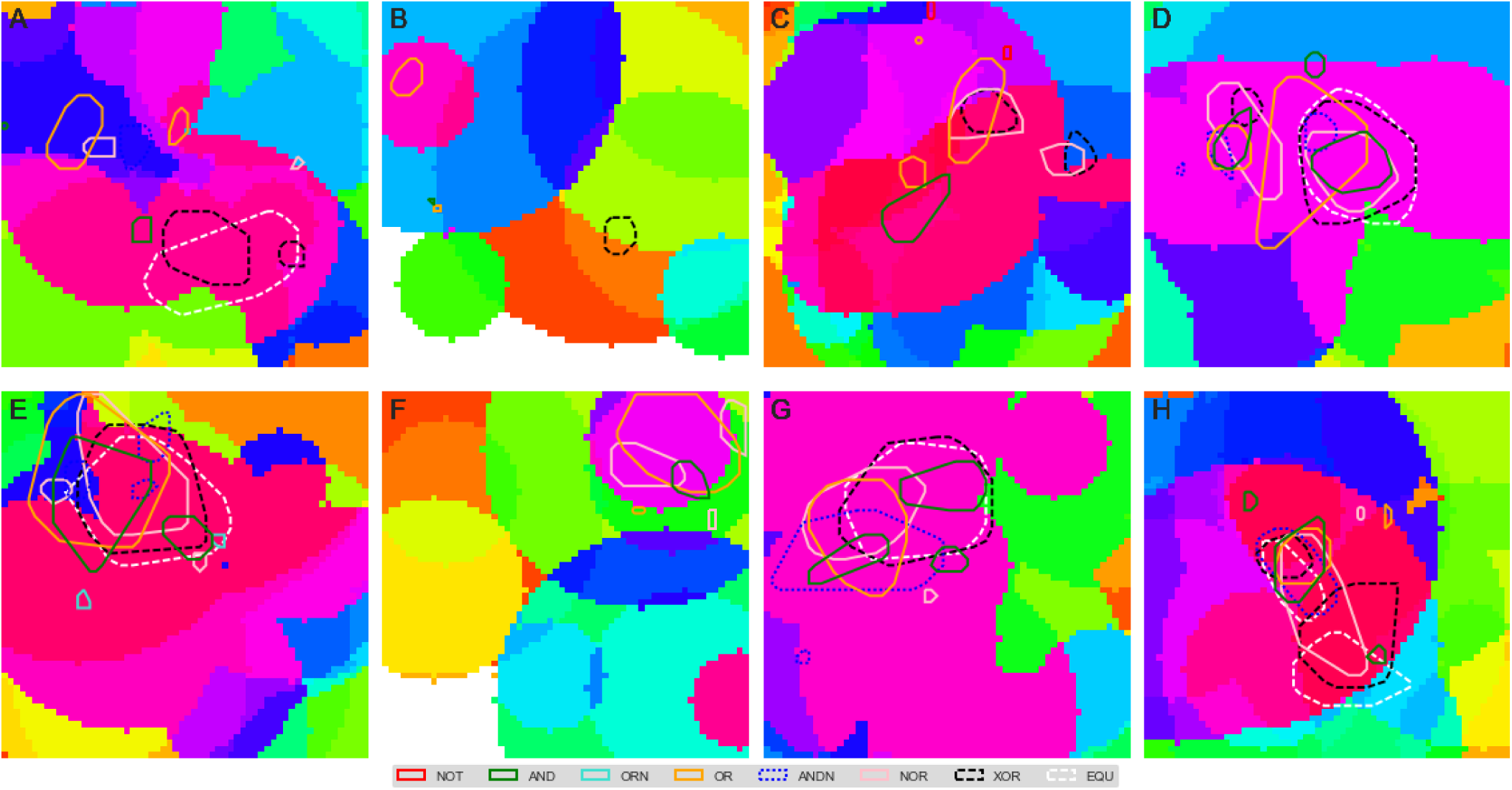
Hotspots across all traits and all environments. Each panel represents a different environment. Background colors represent the different combinations of resources that are present in each location; each color represents a different combination, with colors assigned such that higher hue values roughly correspond to more complex environments (either in number of resources or complexity of traits associated with resources). Each polygon color represents the outline of hotspots for the trait specified in the legend. Note that only hotspots significant at an alpha-level of .00001 are shown. This conservative value was necessary to be certain that hotspots are present despite the large number of hypothesis tests performed. However, in practice, it likely translates to an underestimation of the true size of many of the hotspots.

In most environments, there appears to be strong overlap in hotspots for the evolution of different traits (see Figure 3). However, there are some environments in which this is not the case, and even the environments with the neatest overlap have some traits with non-overlapping hotspots. Some amount of overlap is to be expected, for three reasons: 1) If any trait is easier to evolve in a given region, then lineages living in that region are more likely to be successful as a result of having evolved the trait, giving them an older evolutionary history, 2) many traits serve as building blocks for other traits, so possessing some traits is an indicator that a lineage is in a part of the fitness landscape from which other traits are easier to reach, and 3) some traits are close to each other within the fitness landscape (for example, the two most complex traits, XOR and EQU, which often have hotspots in nearly identical locations). The fact that the overlap is not universal suggests that the local environment has effects on the likelihood of evolving a trait that go above and beyond the ability of lineages to perform other traits.

### Potential drivers of hotspots

Now that we have established that a spatial pattern exists in the locations where traits first evolve, the obvious next question is: what causes this pattern? Since all traits have potential to serve as building blocks for each other, the simplest explanation would be that locations with more resources have higher evolutionary potential. Note that because we are considering only the first mutation to confer each trait, the resource for that trait cannot exert selective pressure.

To test this explanation, we performed a multi-variate linear regression attempting to predict the value of each position in the kernel intensity map from a series of boolean variables indicating which resources were present in a cell. We tried such regressions on various subsets of the data: a single trait within a single environment, a single trait across environments, and all traits across all environments. Models built from data on a single trait in a single environment varied wildly in the percent of variation that they explained, but many explained a high percentage (*R*^2^ values ranged from .11 to .95). However, little (if any) of this explanatory power generalizes across environments. Resources that increase evolutionary potential for a given trait in one environment (as indicated by a regression coefficient significantly greater than 0) may decrease it in a different environment (as indication by a regression coefficient significantly less than 0). Linear models calculated across environments were unable to reconcile these differing patterns, explaining minimal variation in evolutionary potential (*R*^2^ values ranged from .03 to .13). A similar set of models in which the predictor variables were the distance to each resource was similarly ineffective. While high degrees of spatial autocorrelation in the kernel intensity maps mean that the assumption of independence is violated for these linear models, such a reduction in degrees of freedom should only inflate the apparent variation explained. The failure of this simple mechanism to explain our observed results suggests that more complex processes are driving these effects. Note: Interestingly, there does appear to be a correlation between hotspot location and the pink background color in figures 3 and 5. This suggests that the color selection algorithm may be making useful generalizations that are worth further exploration. Nevertheless, it does not appear to be a complete explanation (not all pink regions are hotspots and not all hotspots are in pink regions) and further exploration will be necessary to determine what factors are driving the correlation.

Many hotspots appear to overlap with borders between different combinations of resources. Such regions likely have elevated local diversity, as different phenotypes will be better adapted to the different regions. As discussed above (see Background), there is reason to believe that diversity leads to increased evolutionary potential. Thus, variation in local diversity is another possible driver of evolutionary hotspots. To test this hypothesis, we calculated local diversity across the environment for time points immediately before a novel trait first appeared. Specifically, we measured local diversity as the Shannon entropy of phenotypes within the 25 cells in the 5x5 patch around a given focal cell. We performed this calculation for each cell in the environment at a given time point, producing a map of local diversities. Within each of these maps, we calculated a percentile for each of the 3600 local diversity values, based on the proportion of sites that had a lower diversity.

Next, we located the cell in which a novel trait was about to evolve and recorded its percentile. If diversity has no impact on the probability of a new trait evolving, we would expect all percentile values to be equally likely (i.e. follow a uniform distribution). If, on the other hand, new traits are more likely to evolve in areas with elevated local diversity, we would expect high percentiles to be overrepresented.

To test this hypothesis, we compared the distribution of observed percentiles to a uniform distribution. For the simplest traits, low percentiles were overrepresented due to the presence of populations that contained large swaths of area populated by minimally evolved organisms with none of the traits. However, among the five most complex traits, we found no substantial deviation from a uniform distribution (see Figure 4). This result suggests that traits did not evolve more frequently in more diverse regions. While it is possible that this lack of an effect is the result of the specific diversity metric used, it strongly suggests that local diversity is not an important driver of hotspot location. This result may also cast doubt on the idea that hotspots of diversity are hotspots of all evolutionary processes, but we would need to examine phylogenetic diversity to fully address that point.

**Figure 4:**
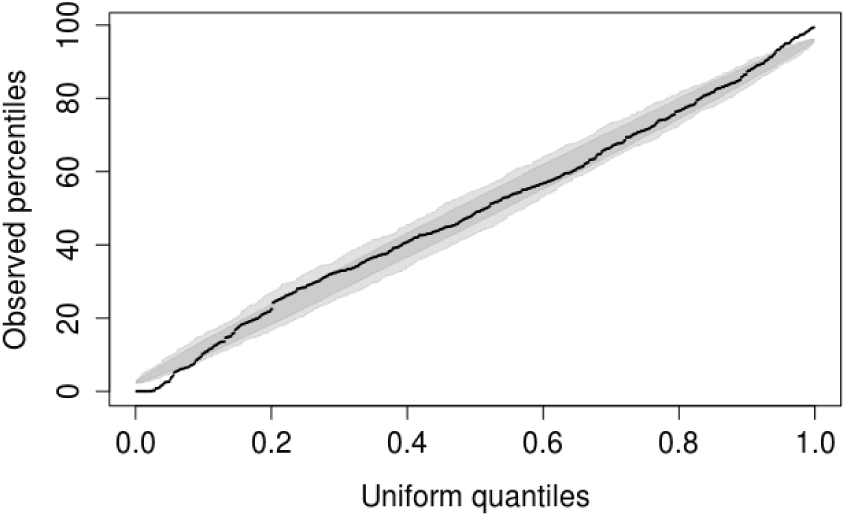
Quantile-quantile plot comparing the distribution observed diversity percentiles for the most complex trait (EQU) to a uniform distribution. (Oldford, 2016). The shaded area represents the 95% confidence interval for where we would expect this line to fall if these distributions were identical. Since the line is mostly within this area, we conclude that the distribution of ranks does not deviate substantially from a uniform distribution. This finding suggests that there is nothing special about the level of diversity in the regions where traits are more likely to evolve. Results for other complex traits are similar.

A third potential driver of evolutionary hotspots is the temporal series of local conditions that lineages experience over evolutionary time as each offspring is born in a slightly different location than its parent. Perhaps some sequences of environments are more conducive to the evolution of certain traits than others. To explore this idea, we located the first organism with each trait across all replicates and traced the spatial path of their lineages. We then overlaid these paths on the corresponding environment (see Figure 5). While a thorough statistical analysis of the variation in paths is beyond the scope of this paper, we do see some qualitative patterns. Indeed, some traits display consistent patterns that are not seen in paths generated from the control data. These patterns include the middle portions of paths, indicating that not just the ending conditions are likely to matter. For instance, in the three environments in which most of the hotspots are overlapping and in the center (D, E, and G in Figure 5), it is rare for any lineage that has ever gone through the lower right corner to be the first one to evolve the most complex task (EQU). This pattern is almost certainly related to the fact that all replicates are seeded with a single organism in the upper left corner; in these environments, successful lineages go directly to the hotspots and don’t leave. In other environments, however, successful lineages appear to travel through different sequences of regions (see Figure 5F, as an example). While more research is required to understand the mechanisms behind such patterns, this effect is the most promising explanation for the existence of evolutionary hotspots.

**Figure 5:**
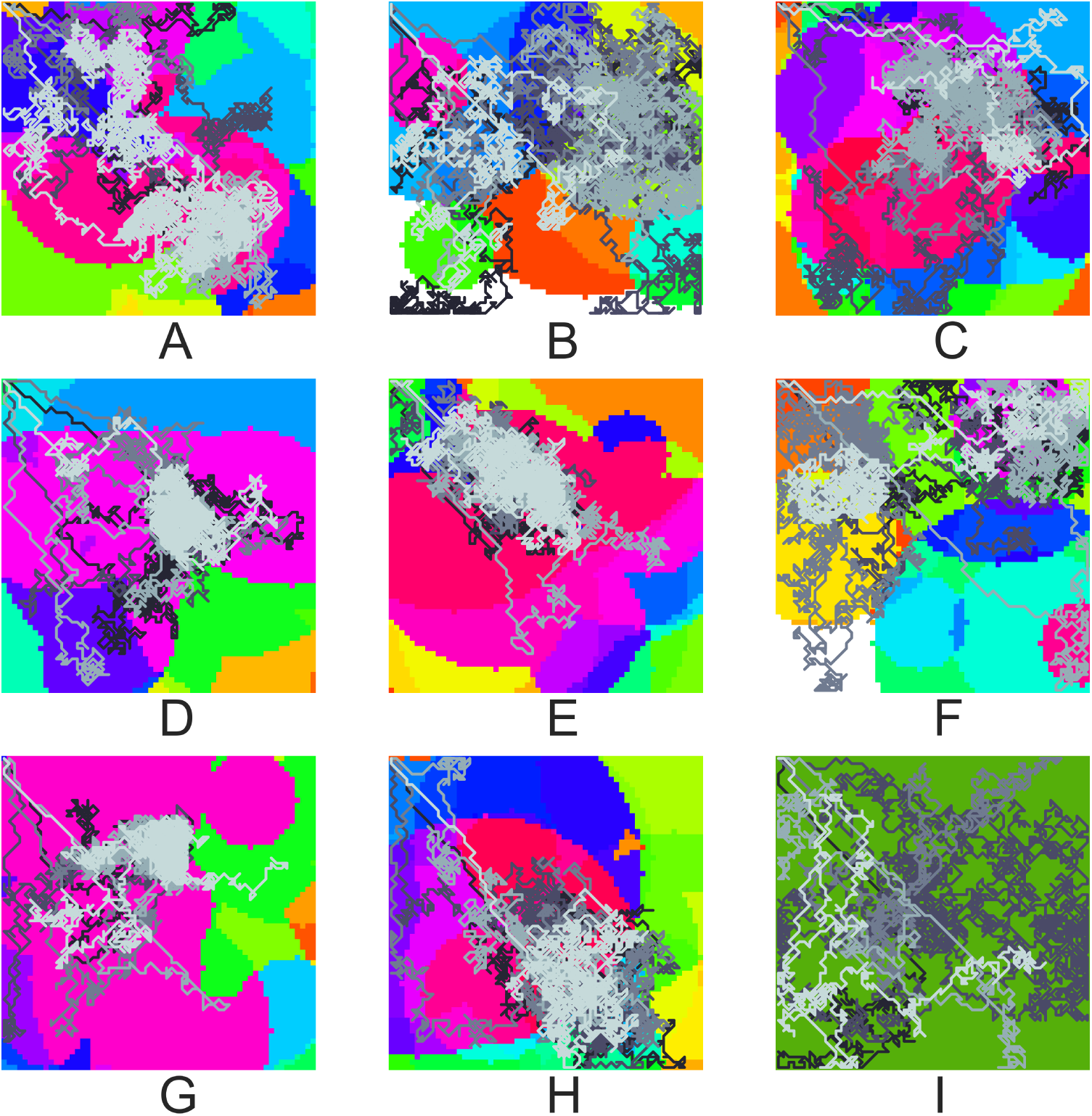
Spatial paths taken by the first lineages to evolve the most complex trait (EQU) from five arbitrarily-selected replicates from each environment. The path from each replicate is colored a different shade of gray. As in Figure 3, background colors represent the set of resources in each part of the environment. I) shows the homogeneous control environment.

## Conclusions

We have demonstrated the existence of spatial variation in evolutionary potential across a heterogeneous environment. Research on what specific aspects of a local region are responsible for elevated evolutionary potential is ongoing. The failure of obvious explanations, such as resource presence/absence and local diversity, to predict the location of hotspots suggests that the actual drivers are complex and nuanced. Preliminary investigations of the sequence of locations that a lineage passes through suggest that this avenue is promising for further exploration; there appear to be broad commonalities among paths of lineages that evolved the same trait in the same environment.

Understanding the drivers of spatial variation in evolutionary potential would give us a valuable tool for controlling evolutionary trajectories. The most immediate applications are in evolutionary computation. Many evolutionary algorithms vary the fitness landscape over time, starting with easy problems and working up to more challenging problems (Hornby, 2006; Ovaska et al., 2009). This approach often requires hand-tuning of the different problems and when they appear in the environment. Varying the fitness landscape across space instead could achieve the same outcome. If we can purposefully build spatial networks of fitness landscapes to promote the evolution of a solution to the problem, that would make such an approach even more powerful.

In the longer term, once our findings have been replicated in a biological system, we hope to use these same concepts to promote the evolution of biological traits of interest. For example, we may be able to engineer habitats that increase the chances of species adapting to climate change through evolutionary rescue. Conversely, we may also be able to arrange environments to inhibit the evolution of undesirable traits, such as antibiotic resistance in bacteria. Overall, this line of research presents a variety of promising applications.

## Acknowledgements

We extend our thanks to Lindsay Williams, Ashton Shortridge, Michael Wiser, Alexander Lalejini, and Rosangela Canino-Koning for suggestions on the manuscript. This research has been supported by the National Science Foundation (NSF) BEACON Center under Cooperative Agreement DBI-0939454, by an NSF Graduate Research Fellowship to ED under Grant No. DGE-1424871, and by Michigan State University through computational resources provided by the Institute for Cyber-Enabled Research. Any opinions, findings, and conclusions or recommendations expressed in this material are those of the author(s) and do not necessarily reflect the views of the NSF.

